# Pelagic trophodynamics control invertebrate population dynamics

**DOI:** 10.1101/2025.11.20.689514

**Authors:** Leocadio Blanco-Bercial, Fernando González Taboada, Joanna M. Pitt, Jirani L Welch, Tammy M. Warren

**Author notes:** **Materials & Correspondence** Leocadio Blanco-Bercial. Bermuda Institute of Ocean Sciences – Arizona State University. 17 Biological Station St. George’s, GE 01, Bermuda.

## Abstract

Invertebrates are increasingly targeted by commercial fisheries, yet assessment approaches designed for finfish often fail to capture their complex population dynamics. For many crustaceans, prolonged, long-distance larval dispersal decouples local recruitment from adult abundance and exposes populations to remote climate conditions overlooked in traditional models. Here we introduce a trophodynamic index that anticipates adult invertebrate population fluctuations by tracking material flows through the pelagic food web during the dispersal and early growth period. Combining primary production with the slope of zooplankton size spectra, the index reflects both productivity and trophic transfer efficiency. Applied to Bermuda’s Caribbean spiny lobster population, which supports both commercial and recreational fisheries that have steadily declined over the last decade, ecological forecasts based on this trophodynamic index outperform climatological, persistence, and environment-based models. This framework links pelagic ecosystem processes to invertebrate population dynamics, improving predictive capacity and supporting the development of ecosystem-based management for small-scale fisheries.

## Introduction

Invertebrates are increasingly targeted by commercial fisheries ^1, 2^, yet the assessment and management of these fisheries challenges approaches developed for finfish fisheries, especially in the context of ecosystem-based management. In the case of crustaceans, such as the spiny lobster, intricate life cycles induce complex, supply-side dynamics ^3^. Long pelagic dispersal phases, for example, decouple density-dependent recruitment from the local spawning stock and expose the population to remote climate conditions ignored in traditional assessments^4^.

The Caribbean Spiny Lobster, *Panulirus argus* (Latreille, 1804) is a spiny lobster (Family Palinuridae) whose adults can be found between 35 N and 25 S of the coastal Western Atlantic, with occasional records off the coast of Africa^5^. This species has been heavily exploited across its range ^6^, and it is widely assumed that weak regulatory frameworks, combined with overfishing, have contributed to a general decline in population abundance throughout its distribution^6, 7, 8, 9^.

The life cycle of *Panulirus argus* in Bermuda features spring mating and summer brooding of the fertilized eggs, with hatching larvae being released into the water column near the edge of the shallow platform in late summer. The phyllosoma larvae spend several months in the upper ocean as part of the planktonic community, feeding first on gelatinous zooplankton and later on larger pelagic crustaceans^10^. Over this period, the larvae molt eleven times before metamorphosing into the puerulus postlarval stage as they approach coastal waters^11^. In Bermuda, this stage settles to the seafloor in late summer^12^. Early benthic juveniles are solitary, inhabiting shallow algae and seagrass areas, but later aggregate and migrate to deeper waters, where they continue to molt and grow into adults.

The long pelagic stage of *P. argus* causes the populations to be admixed, and local recruitment might not be the rule for most locations ^13, 14^. Near Bermuda, prevailing surface currents make the return home very unlikely for locally produced larvae but favor the arrival of larvae from the west. The population is thus considered a sink for Caribbean larvae, but also a source of propagules that drift further north and east, providing a steppingstone to colonize new habitats. As a consequence, recruitment in Bermuda mostly depends on the supply of larvae from Caribbean populations, and it is decoupled to a large extent from local adult abundance.

The Bermuda spiny lobster fishery is a small-scale, coastal fishery operating seasonally from September to March, across both inshore (<10 m) and offshore (10–100 m) grounds.

Following a severe decline in local finfish stocks in the 1980s, the traditional arrowhead traps that were used to target both fish and lobsters were banned in 1990. However, the lobster fishery was doing well at that time, so a small, experimental fishery to develop a lobster-specific trap took place from 1991 to 1996 ^15^, and a limited-entry fishery utilizing these new, standardized traps began in September 1996. Over the years, between 18 and 31 fishers deploying 8 to15 traps per license have operated each season, distributed between the western and eastern ends of the island. Management measures regulate the level and duration of trapping in inshore areas (inside the 10 m isobath; Figure 1) compared to the offshore (10 to 100 m. (Supplementary Table S1). The trap design, specifically the entrance funnel and escape gap dimensions, selects for lobsters aged approximately four to five years, thereby excluding juveniles and protecting large, highly fecund females. The Department of Environment and Natural Resources (DENR) also regulates the recreational fishery in which free divers use poles with nooses to capture lobsters. The number of recreational licenses, first capped at 500 in 2017, was reduced to 340 in 2022 and to 175 in both 2023 and 2024. Catches from the recreational sector typically account for 10% of the commercial harvest.

**Figure 1.**
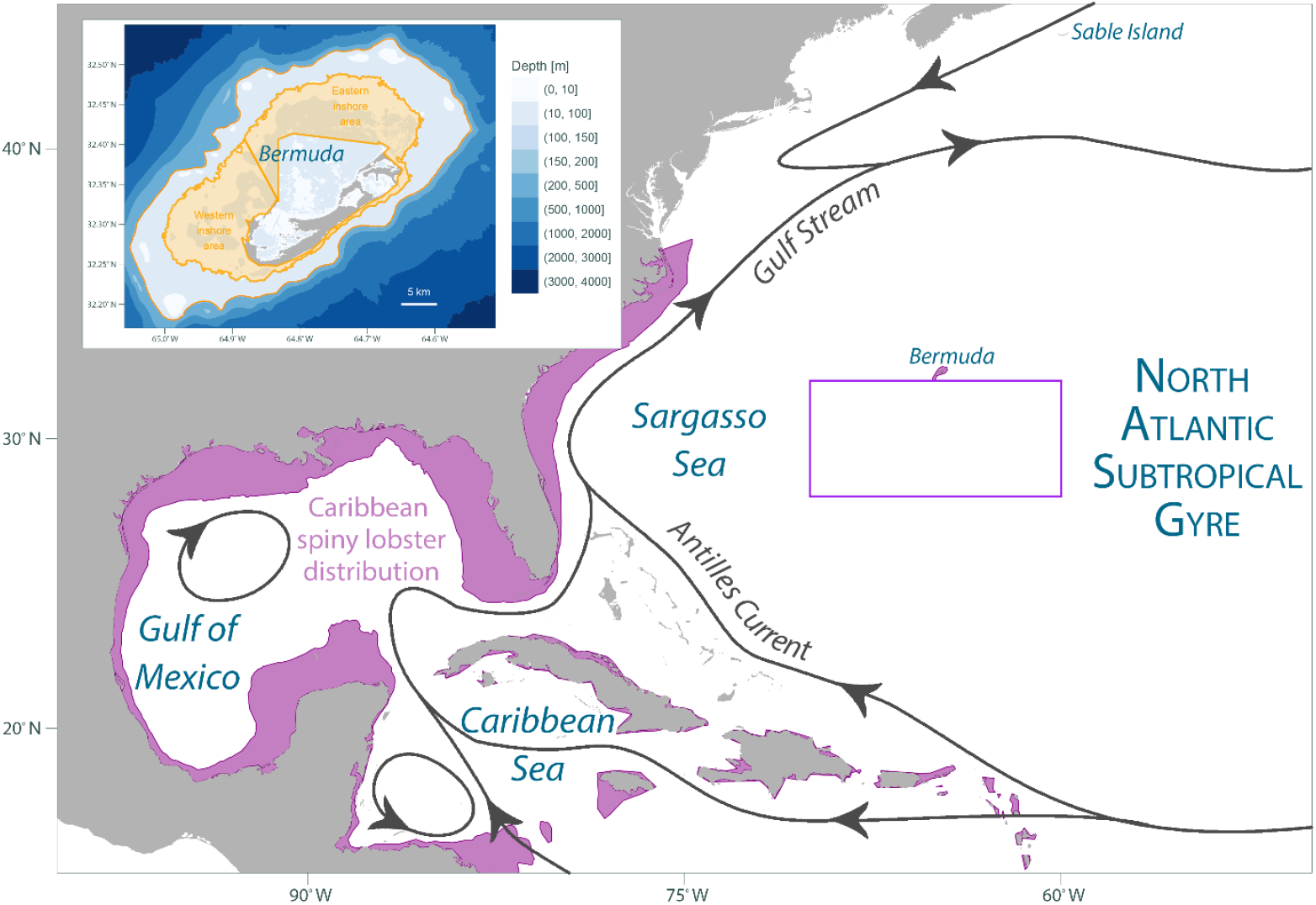
Partial distribution of the Caribbean Spiny Lobster, and inshore and offshore fields of the Bermuda fishery. The square box was used to calculate regional trends in environmental conditions (see methods for details).

To overcome the uncertainty that results from the decoupling of local spawning stock and recruitment, in this manuscript we introduce a trophodynamic index to anticipate fluctuations in adults based on material flows through the pelagic food web over the larval dispersal and growth period:

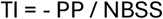

The index combines regional estimates of primary production (NPP) and the slope of the zooplankton size spectra (NBSS) to capture fluctuations in overall productivity and tropic transfer efficiency.

We illustrate the approach with the Caribbean spiny lobster population in Bermuda, which supports both commercial and recreational fisheries that have steadily declined to historic lows over the last decade. This index, based on the trophodynamic dynamics, beat forecasts based on climatological levels and on local persistence, as well as forecasts incorporating physical or biogeochemical variables such as temperature and primary production. The trophodynamic index bridges a gap to improve the management of small-scale invertebrate fisheries and paves the way for the development of more complex, predictive, ecosystem-based management approaches

## Results

### Environmental variability in the subtropical North Atlantic

To characterize changes in environmental conditions affecting the dynamics of the Caribbean spiny lobster in Bermuda, we analyzed changes in surface temperature, and the components of the proposed trophodynamic index; satellite net primary production (NPP) and the slope of the normalized biomass size spectra (NBSS) of zooplankton. All these variables show trends consistent with those previously reported. Temperature averaged over the selected area (Figure 1) shows an increase after 2010, while NPP decreased (Figure 2). The NBSS data follow the patterns discussed by Russo et al. ^16^, although there are minor differences due to the September to August yearly averages compared to the natural year averages. The trophodynamic index derived from the ratio between NPP and NBSS (average between day and night NBSS) showed a maximum peak for 2010, decreasing afterwards (Figure 2).

**Figure 2.**
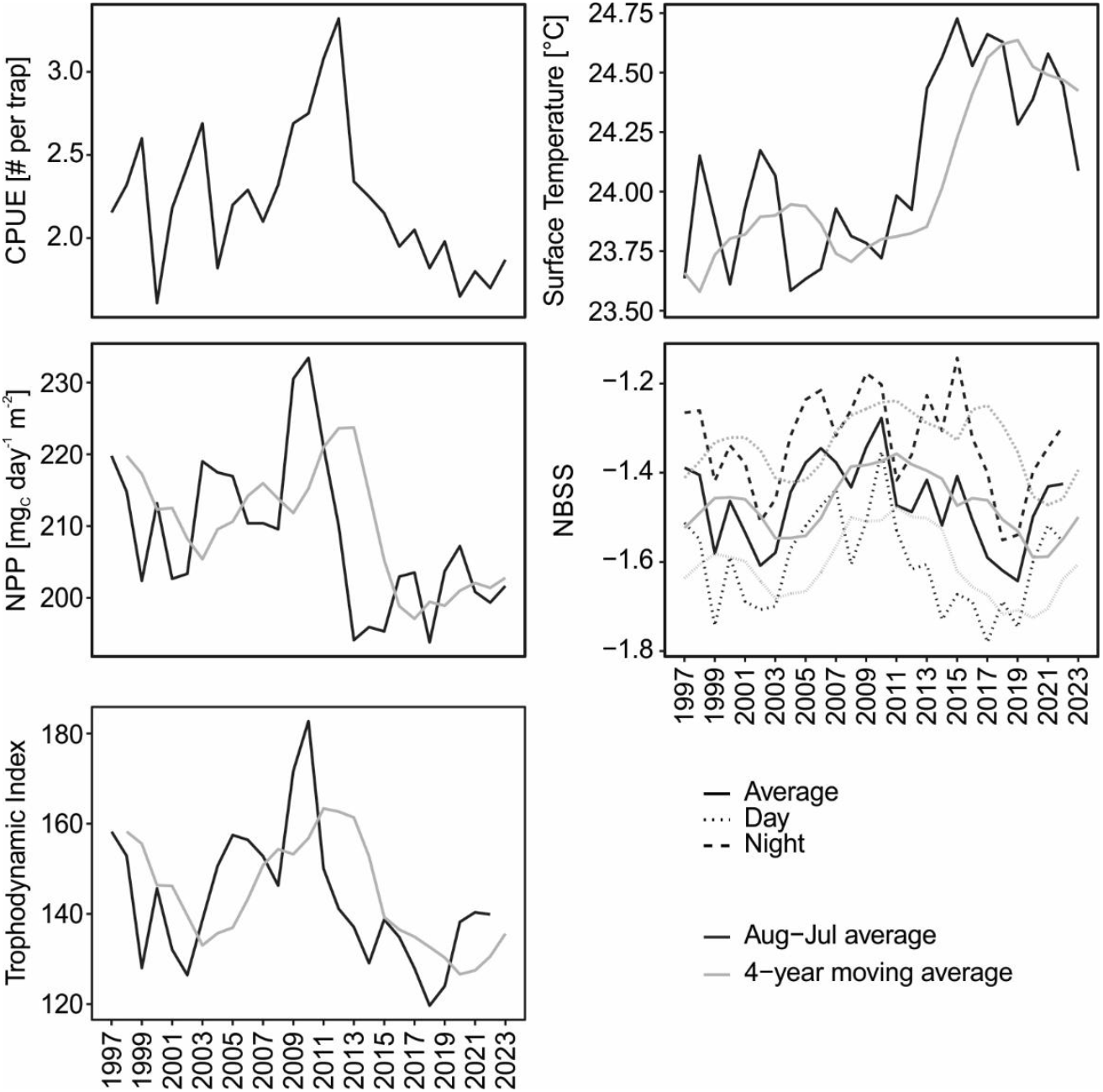
CPUE, satellite surface temperature of the ocean, satellite-derived net primary production (NPP), NBSS, and Trophodynamic Index series 1997 to 2023.

### Fishery data

After reopening in 1997, the lobster fisheries underwent two distinct phases (Figure 2). After some initial variability, total catch per season rose from 19,147 individuals in 2000 to a peak of 37,323 in 2011. Then, total catch steadily declined down to 8,095 in 2024 (with a record low of 7,061 lobsters in the 2023 season, an interannual decline of 10.4%). Accounting for the intensity of fishing activity by standardizing against the number of traps hauled, the average catch per unit effort (CPUE) has oscillated between 1.61 and 3.32. The general trend showed a maximum peak during 2011-2013, followed by a steady decline (Figure 2).

### Statistical Analyses, Forecast and Evaluation

High variability of spiny lobster CPUE precluded anticipating fluctuations from the climatological average of observed past values^17^ (CV = 45%, RMSD = 0.46; Table 1 and Figure 3). However, forecasts based on the persistence of previous year’s CPUE rendered some success, establishing a considerable benchmark for forecasts based on environmental variables to surpass (r = 0.68, RMSD = 0.36).

**Table 1.** Statistical metrics assessing the skill of forecast models informed by temporal changes in the trophodynamic index, net primary production (NPP), and temperature at the ocean surface (TOS), as well as two additional benchmark forecasts based on the average CPUE from all years prior to the forecast year (climatology), and the CPUE from the immediate previous year (persistence).

| Predictor | Mean absolute scaled error, MASE |  |  | Pearson correlation, *r* |  |  | Root mean square deviation, RMSD |  |  |
| --- | --- | --- | --- | --- | --- | --- | --- | --- | --- |
|  | Target | Model | Potential | Target | Model | Potential | Target | Model | Potential |
| **Index** | 0 | 0.86 | **0.68** | 1 | 0.73 | **0.87** | 0 | 0.31 | **0.22** |
| NPP | 0 | 1.25 | 1.05 | 1 | 0.29 | 0.53 | 0 | 0.42 | 0.37 |
| TOS | 0 | 0.95 | 0.84 | 1 | 0.64 | 0.70 | 0 | 0.36 | 0.32 |
| Climatology | 0 | 1.45 |  | 1 | 0.00 |  | 0 | 0.46 |  |
| Persistence | 0 |  | 1.00 | 1 |  | 0.68 | 0 |  | 0.36 |

**Figure 3.**
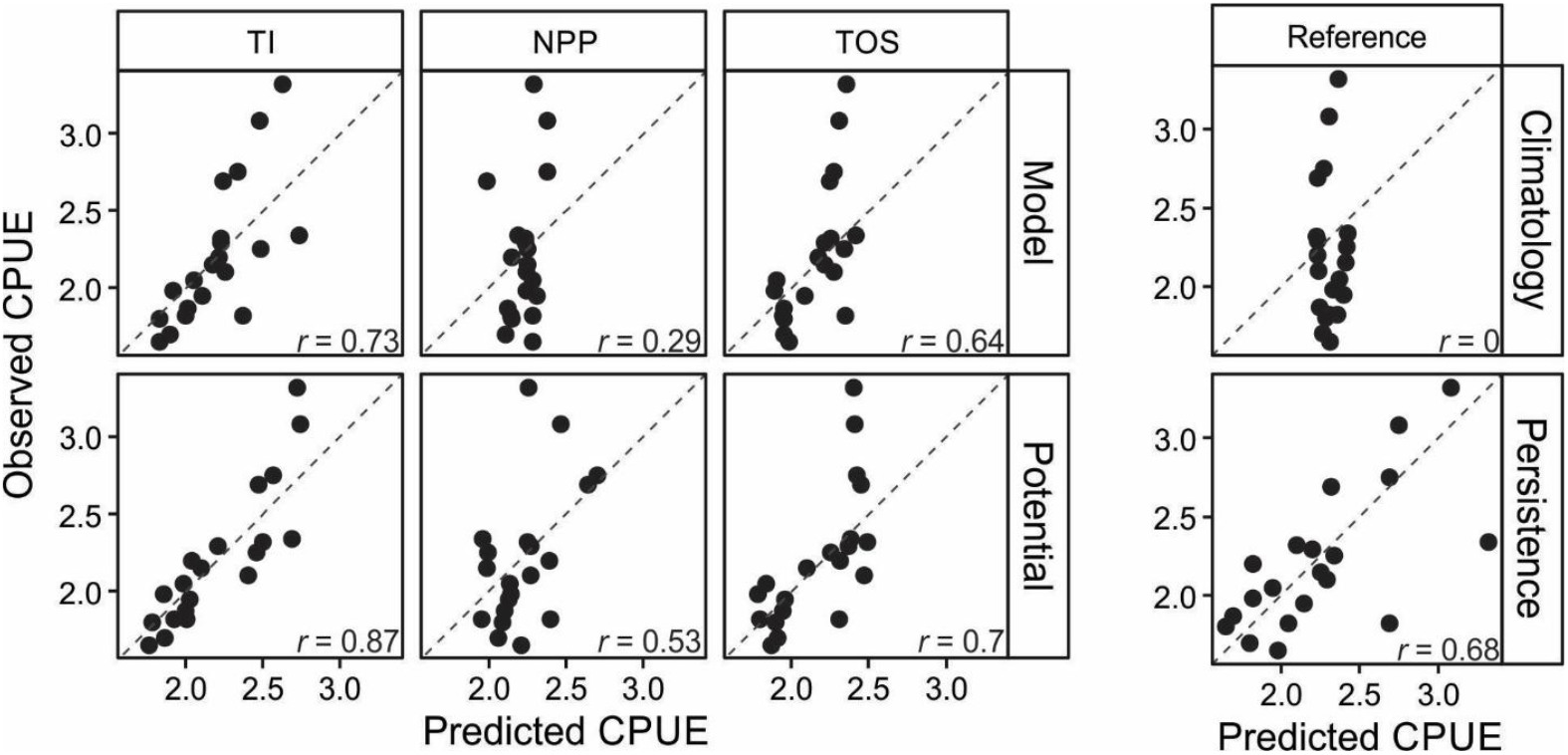
Correlations between predicted vs observed CPUE, based on the regressions derived from the Trophodynamic Index (TI), the satellite-derived Net Primary Production (NPP) and temperature at the ocean surface (TOS), as well as based on Climatology and Persistence (see methods). The best correlations were obtained for the TI, both for the model and the potential calculations.

These models used a moving average to incorporate variability in environmental conditions over the previous 4 years to match the prolonged larval and juvenile period of the Caribbean spiny lobster. The fit between the CPUE and the previous 4-year moving average trophodynamic index (R^2^ = 0.50) was higher than that of TOS (R^2^ = 0.34) and PP (R^2^ = 0.19; Figure S1 in Supplementary file 2). Besides, the forecast based on the trophodynamic index was the only model able to overperform the persistence forecast in all three metrics considered (Figure 3; Table 1). All these figures were robust to considering either potential estimates based on all available observations, or the more realistic figures above based on an iterative assessment mimicking the growing availability of observations in time (Potential *vs* Model in Figure 3 and Table 1). The variance explained by forecasts based on the trophodynamic index decreased by only 30% when comparing realized *vs* potential forecasts, which suggests robust estimates of the effect even in short time series. This value was intermediate between the 17% decrease observed for forecasts based on seafloor temperature and the 70% decline recorded for forecasts based on primary production.

## Discussion

We demonstrated the potential for anticipating fluctuations in the abundance of the Bermuda population of the Caribbean spiny lobster, a marine invertebrate with an open population, based on simple trophodynamic principles. To this end, we developed a new index based on ocean primary production and the efficiency of zooplankton in transferring energy to upper size classes. Forecasts informed by this simple trophodynamic index outperformed forecasts based on climatology or persistence, or incorporating information from other relevant environmental variables.

The index relies on satellite observations (NPP) and time-series data (zooplankton normalized biomass size spectra, NBSS) from the Bermuda Atlantic Times Series (BATS), both of which are public resources and independent of fisheries data and reports, two desirable characteristics for forecasting tools ^18^. While the decline in ocean primary production in the region where these lobster larvae spend their planktonic phase is the strongest in the world ^19^, NPP alone did not adequately explain decline in the abundance of spiny lobster. Our index captures not only the decline in NPP, but also the consequences of this decline and the changes in identity of the producers on the zooplankton community, as represented by the NBSS.

The trophodynamic index was able to capture two of the main effects of the ongoing oligotrophication of the oceans. Increased ocean stratification reduces nutrient availability in the epipelagic zone, leading to lower primary productivity ^19, 20^, and also causes a shift toward a community dominated by smaller-sized primary producers ^21, 22^. This shift also results in less efficient trophic transfer by zooplankton to higher trophic levels ^16^. Together, these effects magnify the impact of oligotrophication on fisheries beyond what could be explained by net primary production alone ^23, 24, 25^.

Previous efforts have relied on direct measurements of bulk ecosystem properties such as Chl-a or NPP ^18^, or on complex models that incorporate, among other variables, NPP along with trophic transfer ^23, 26^. Our index targets a midpoint by combining the simplicity of the former and the ecological realism of the latter through basic trophodynamic principles. Other approaches relied on pelagic NBSS to calculate the percentage of phytoplankton biomass that would be transferred to upper trophic levels as a proxy for maximum sustainable fish biomass ^25^. Our approach attempts to capture the dynamic nature of planktonic production and the rapid consumption by zooplankton through the combination of production rates and transfer efficiency instead of focusing on the standing stocks of both producers and the pelagic system.

Another advantage of the proposed index is that it relies on variables often available from plankton monitoring. Primary production is routinely measured in the field and estimated using remotely sensed data. The slope of the size spectra can be estimated from size-fractionated zooplankton biomass datasets, which are often part of standard time-series measurements e.g., ^27^. These types of measurements are expected to become more widely available with the widespread implementation of imaging technologies e.g., ^28, 29^. Oceanographic satellites have recently started to provide estimates of zooplankton standing stocks ^30^, and are increasingly focusing on the retrieval of plankton functional groups and size structure ^31^.

Although lobsters are no longer part of the plankton community once they have settled, reef processes and productivity in Bermuda are strongly influenced (and sustained) by offshore dynamics and primary production ^32, 33^, including zooplankton that support coral heterotrophy and the pelagic primary production that sustains benthic life in the intermediate depths around the edge of the Bermuda platform. This index, though based on oceanic variables, would therefore reflect similar trends occurring on the Bermuda platform which could continue to impact the early benthic survival of juvenile Caribbean spiny lobsters, before entering the fishery.

One missing piece of the puzzle regarding the lobster fishery stock is the contribution of Bermuda’s local population to recruitment. Due to the species’ long planktonic phase, it is highly unlikely that Bermuda lobsters significantly contribute to self-recruitment. Previous genetic studies have found significant differentiation (*F*_ST_) only with populations from Panama ^34^, suggesting a quasi-panmictic population across the species’ distribution. Moreover, due to the absence of a return current, Bermuda lobsters may contribute minimally to recruitment—either locally or to the broader Caribbean ^13^.

In sum, this index, which is independent of both fishery effort and stock assessments, fills a critical gap in improving the management of this small-scale invertebrate fishery and paves the way for the development of more complex, ecosystem-based management approaches.

## Methods

Scripts for data collection, data, and calculations, are available at https://gitlab.com/biosasu/lobster.

### Environmental data

We relied on reanalysis data to reconstruct the environmental conditions experienced by Bermuda spiny lobster across relevant temporal and spatial scales, including the conditions of the open ocean. Considering the location of Bermuda in the North Atlantic, the circulation of the subtropical gyre, and the long duration of the phyllosoma stage, we retrieved and analyzed data for a broader region beyond the coastal Bermuda area. Sea water temperature at surface (TOS) was obtained from the GLORYS12V1 hindcast, which is based on Copernicus’ Global Ocean Physics Reanalysis ^35^. Daily values of primary production were retrieved from Copernicus Global Ocean Biogeochemistry Hindcast ^36^. To reduce potential mesoscale and local effects — and to better reflect regional trends — data were averaged over the area bounded by 28° to 34° N and 60° to 70° W (Figure 1). From the daily values, seasonal averages were calculated for cycles spanning September 1 to August 31, aligning with both the reproductive cycle (i.e., starting the yearly cycle immediately after spawning) and the fisheries calendar. Reanalysis data were available from 1997 onward.

### Zooplankton Normalized Biomass Size Spectra

Data from the Bermuda Atlantic Time-series Study site (BATS), available at https://www.bco-dmo.org/project/2124 (DOI: 10.26008/1912/bco-dmo.881861.5) ^21, 37^ were used to calculate the zooplankton Normalized Biomass Size Spectra (NBSS) ^38^. Following Mehner et al. ^38^ and Russo et al. ^16^, size-fractionated zooplankton biomass from the upper 200 m of the water column was converted to carbon biomass using the equations from Madin et al. ^39^: for day and night samples (C_biomass_ = 0.36 * Biomass for day samples, C_biomass_ = 0.37 * Biomass for night samples). For each data point, average values were taken for each size fraction, based on two replicates per cruise for both day and night sampling events. NBSS was then calculated as the slope of the regression between the Log_10_-transformed biomass (normalized by bin width) and the midpoint of the Log_10_-transformed size bins for each fraction.

Size fractions were defined as 200–500 µm, 500–1000 µm, 1000–2000 µm, 2000–5000 µm, and 5000–30,000 µm. Although zooplankton are typically defined as organisms smaller than 20 mm, our upper limit better reflects the composition of zooplankton samples at BATS (Blanco-Bercial, pers. obs.) and aligns with the approach used by Russo et al. ^16^.

NBSS slope values were calculated separately for day and night samples, and only statistically significant slopes were included in our analysis. We also computed the average between day and night values, as lobster larvae are exposed to both communities during their diel vertical migration. Moreover, the overall trophic transfer efficiency of the system is likely influenced by both day and night conditions.

### Definition of a new index

Trophic efficiency and plankton production have been traditionally invoked to explain large scale spatial gradients in the production of higher trophic levels ^40^. The inability to simultaneously characterize both aspects with enough resolution, however, limits the application of trophodynamic principles in fine scale settings. The NBSS slope provides a useful measure of trophic transfer efficiency within the zooplankton community and toward higher trophic levels ^16, 41, 42, 43^. The NBSS slope, however, only reflects energy transfer efficiency and, according to theory, it can be largely decoupled from changes in absolute energy availability ^42, 44^.

Spatial and temporal gradients in primary production quantify changes in the amount of energy entering the pelagic ecosystem through photosynthesis, and provide a proxy of regional to large-scale variation in fishery productivity ^45, 46, 47^. Changes in primary production, however, need to account for the number and efficiency of links between producers and fish ^24, 45, 46, 47^. A more recent approach uses pelagic NBSS to calculate the percentage of phytoplankton biomass that would be transferred to support fish biomass ^25^, yet both of these metrics represent standing stocks and may fail to reflect the dynamic nature of planktonic production and the rapid consumption by zooplankton.

To capture both the dynamic processes involved in ocean productivity (i.e., rates) and the trophic efficiency of the system, we propose the following trophodynamic index (TI), which exploits the inverse relationship between the slope of the size spectra and trophic efficiency ^48^:

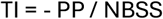

This index is based on a rate rather than a stock, providing a more accurate reflection of available energy and the transfer of ocean productivity. It indicates how much of the primary production (PP) is transferred to upper trophic levels, accounting for both the efficiency of the system and the amount of energy entering it. Higher PP values will lead to higher index values, while steeper NBSS slopes, indicating lower trophic transfer efficiency, will result in lower index values.

### Fishery data

DENR has collected statistics on the Bermuda spiny lobster fishery from 1997 to 2024, including records of the total number of lobsters caught, and the number of traps deployed, thus providing a consistent measure of catch per unit effort (CPUE). Commercial fishers account for the bulk of catches (e.g., 8095 *vs* 965 lobster were caught over the 2022-23 season by commercial and recreational fishers respectively). Data consist of self-reported forms completed by commercial fishers, and these are validated by onboard or shoreside observers for a proportion of the catch. The reporting form for each fishing trip details the number of pots hauled, with associated soak time, and the number of lobsters caught. Based on these data, we estimated catch per unit effort (CPUE) as the ratio between the total number of lobsters caught and the total number of pots hauled, averaged over the entire fishing season. We also assumed that catchability was constant and therefore that changes in our CPUE index were informative about interseasonal changes in total lobster abundance in the area.

### Statistical Analyses, Forecast and Evaluation

To test our hypothesis that the proposed index can forecast lobster fishery catches in Bermuda - outperforming other environmental and physical variables - we conducted regression analyses between CPUE, total captures (as number of lobsters), and various environmental and biological variables and indices. Given the lobster life cycle, in which individuals enter the fishery approximately four years after hatching (spending one year in the planktonic phase in the open ocean and three years in coastal waters), we compared the annual CPUE with the four-year moving average of each environmental variable. This approach accounts for the delayed influence of environmental conditions on fishery recruitment. Individual regressions between the CPUE and each environmental variable were done using a Bayesian generalized linear model via Stan ^49^ in R ^50^.

The performance of our ecological forecast model was evaluated using three benchmark approaches: the average CPUE from all years prior to the forecast year (“climatology”), the CPUE from the immediate previous year (“persistence”), and the model performance using data only up to the year prior to the forecast year (“potential”) ^18, 51^. Performance of each metric and the model was assessed using three statistical measures: the coefficient of determination (R^2^), root mean square deviation (RMSD), and mean absolute scaled error MASE; ^52, 53^. To further assess the performance of our index relative to direct environmental variables, both the model and potential metrics were compared with those derived from primary production and mixed layer depth.

## Acknowledgements

We would like to acknowledge the Bermuda lobster fishermen for providing the data reported in this manuscript, and the NSF Bermuda Atlantic Time-series Study program and personnel (NSF 2241455) for the zooplankton biomass data. LBB acknowledges the support of Simons Foundation International’s BIOS-SCOPE program.

## Author contributions

Conceptualization: LBB. Investigation: LBB, FGT, JMP, JLW, TMW. Formal Analysis: LBB, FGT. Writing – original draft preparation LBB, FGT. Writing - Review & Editing LBB, FGT, JMP, TMW. All authors approved the final version of the manuscript.

## Data availability

Environmental data is available in public repositories (https://www.bco-dmo.org/project/2124). Scripts and lobster abundance data are available in the gitlab repository https://gitlab.com/biosasu/lobster and as a release in BCO-DMO at DOI XXXXX (*to be opened released after acceptance)*.

