## Supplementat Figure 1 for "Pelagic trophodynamics control invertebrate population dynamics"

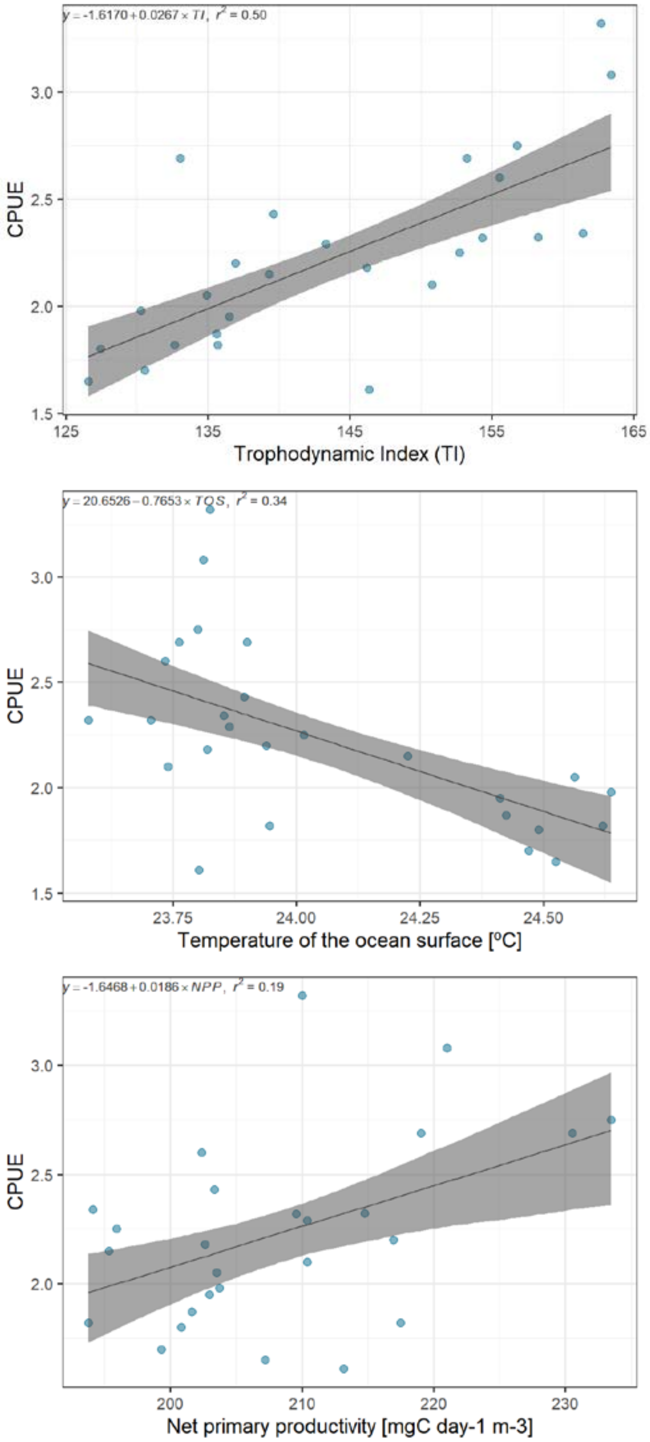


Figure S1. Linear regressions between TI, TOS and NPP with CPUE. The shadowed areas represent the 95% confidence interval.
